# OBMeta: a comprehensive web server to analyze and validate gut microbial features and biomarkers for obesity-associated metabolic diseases

**DOI:** 10.1101/2023.08.07.552363

**Authors:** Cuifang Xu, Jiating Huang, Yongqiang Gao, Weixing Zhao, Yiqi Shen, Feihong Luo, Gang Yu, Feng Zhu, Yan Ni

## Abstract

Gut dysbiosis is closely associated with obesity and related metabolic diseases including type 2 diabetes (T2D) and non-alcoholic fatty liver disease (NAFLD). The gut microbial features and biomarkers have been increasingly investigated in recent studies, which require further validation due to the limited sample size and various confounding factors that may affect microbial compositions. So far, it lacks a comprehensive bioinformatics pipeline providing automated statistical analysis and integrating independent studies for cross validation simultaneously. OBMeta aims to streamline the standard metagenomics data analysis from diversity analysis, comparative analysis, functional analysis, to co-abundance network analysis. In addition, a curated database has been established with a total of 88 public research projects, covering three different phenotypes (Obesity, T2D, and NAFLD) and more than five different intervention strategies (exercise, diet, probiotics, medication, and surgery). With OBMeta, users can not only analyze their own research projects, but also search and match public datasets of interest for cross-project validation. Moreover, OBMeta provides cross-phenotype and cross-intervention-based advanced validation that maximally supports preliminary findings from an individual study. To summarize, OBMeta is a comprehensive web server to analyze and validate gut microbial features and biomarkers for obesity-associated metabolic diseases. OBMeta is freely available at: http://obmeta.met-bioinformatics.cn/.

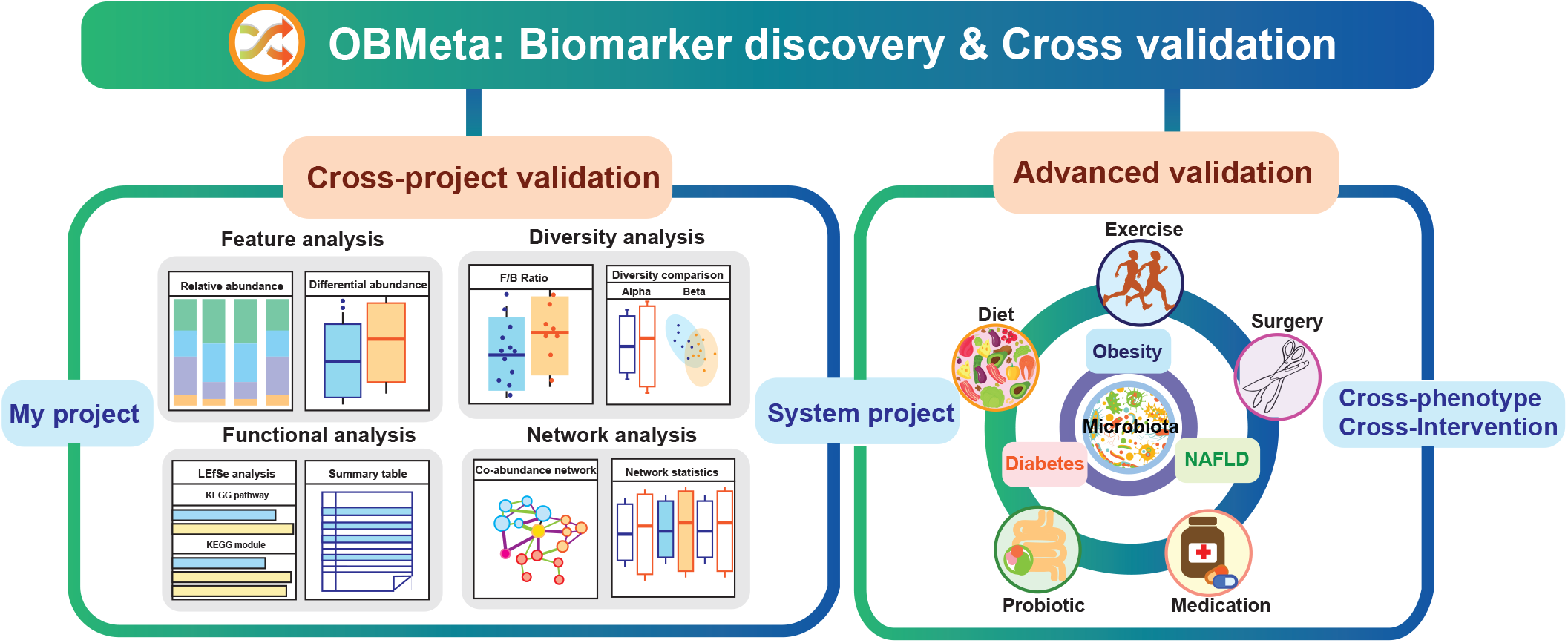

## Key words

Gut microbiome, biomarker discovery, obesity and related metabolic diseases, automated bioinformatics analysis, cross-validation

## Introduction

Obesity is now a global health issue with rapidly increased prevalence that reached 39% worldwide and affected almost 2.5 billion people according to the World Health Organization (WHO) report in 2016 [1]. Previous studies have showed that obesity increases the risk of other metabolic diseases, including type 2 diabetes (T2D) [2], non-alcoholic fatty liver disease (NAFLD) [3] and metabolic syndrome (encompassing hypertension, dyslipidemia, and insulin resistance) [4, 5]. The epidemic of these global obesity-associated diseases causes extensive social and economic implications, so effective intervention is necessary.

Gut microbiota is a complex community of microorganisms inhabiting the host gastrointestinal tract that has crucial roles in host health maintenance [6]. Increasing evidence has demonstrated that dysbiosis of gut microbiota propels the emergence of obesity and correlates with other metabolic diseases. For example, lipopolysaccharides (LPS) derived from pathogenic bacterial membranes can trigger chronic inflammation in T2D and obesity [7, 8]. The proliferation of the short-chain fatty acids-producing bacteria benefits weight loss and enhances insulin sensitivity [9, 10]. In addition, different kinds of weight loss strategies, such as exercise, medication, probiotics intervention [11], vegetarian diet [12], and even stomach surgery [13], can reshape gut microbiota. Therefore, identifying reliable and consistent microbial biomarkers is essential helping us to explore prospective therapeutic targets for obesity-associated metabolic diseases.

With the widespread application of next-generation sequencing technologies in the biomedical field, accumulating number of obesity-associated datasets of gut microbiome has been published in public databases. Metagenomic analyses are challenging due to the large volume and complex nature of raw sequencing data. For non-bioinformatics experts, it is demanding to apply computational programming and algorithms for high-throughput data processing and analysis. Recently, to make more available of comparative metagenomics analysis for clinicians and bench researchers, several powerful web servers have been developed, such as MG-RAST [14], METAGEN-assit [15], MicrobiomeAnalyst [16], and Busybee [17]. They offer efficient microbial data processing, analysis and visualization.

After initial biomarker discovery, further validation is important due to the limited sample size in an individual study and various confounding factors that may affect findings. Since it is time/cost-consuming to conduct independent studies, applying bioinformatics techniques to apply publicly available metagenomic datasets could help us to validate reliable biomarkers efficiently. So far, it lacks a comprehensive bioinformatics pipeline providing automated statistical analysis and cross-project validation with comparable studies simultaneously. To identify and validate microbial biomarkers of obesity associated metabolic diseases, we have developed a comprehensive platform named OBMeta, which implements a complete pipeline for comparative taxonomic, functional profiling and co-abundance network construction. In addition, OBMeta provides cross-project, cross-phenotype, and cross-intervention validation based on the curated and continually updated database. It helps users to identify the gut microbial features and biomarkers of obesity-related diseases efficiently and accurately by evaluating their consistent variations among different studies.

## Materials and methods

### Data collection and curation

Raw sequencing reads (including 16S amplicon raw reads and metagenomic raw reads) were downloaded from NCBI SRA (Sequence Read Archive) [18] and EBI ENA (European Nucleotide Archive) [19] databases using transfer tool Aspera. Sequencing related metadata and host related metadata were obtained from the databases and the published paper respectively. We manually included the datasets based on the following criteria: datasets collected from biological samples (including feces, intestine, oral, and skin etc.) of humans and animals; studies with at least one single-factor comparison on obesity-associated metabolic diseases (i.e., obesity, T2D, or NAFLD); raw data can be linked to at least one published paper. **Supplementary Figure 1** illustrates the detailed pipeline for constructing the curated database of OBMeta.

### Processing of raw data

We evaluated the overall quality of the download data using FastQC (version 0.11.8) and then processed 16S amplicon and metagenomic raw data respectively. **Supplementary Table 1** summarized the software used in the data processing.

For 16S sequences, QIIME 2 [20] and DADA 2 [21] were applied for processing sequences and denoising respectively. The output from DADA 2 was clustered into 99% identity using an operational taxonomic unit (OTU) picking protocol against Sliva 132 database. Functional annotations were conducted by PICRUSt2 (version 2.5.0) [22] with default parameters.

For metagenomic sequences, the sequencing vectors and low-quality bases was removed by Trimmomatic (version 0.39) [23]. The clean reads were mapped against host reference genome using Bowtie 2 (version 2.4.1) [24] to filter out the host contamination. The high-quality reads of single sample were first assembled by MEGAHIT (version 1.2.9) [25]. The unused reads of each sample were mix-assembled again. Next, QUAST (version 5.2.0) [26] was applied to evaluate the quality of the assembling. Then, the assembled reads were predicted open reading frames (ORFs) and filtered (< 100 nt) by MetaGeneMark (version 3.38) [27]. The qualified reads were translated into amino acid sequences and clustering into non-redundant contigs with 95% identity and 90% coverage using CD-HIT (version 4.8.1). Finally, Diamond (version 0.9.34) was used to annotate different taxonomies and functional pathways against NR and KEGG databases with an e-value cutoff of 1E-5, respectively. Different taxonomic relative abundance profiles and functional abundance profiles were determined by Salmon (version 1.2.1) [28].

### Consistency evaluation of bacterial variations among different projects

To evaluate the consistency of a significantly varied taxon among different projects, we defined a score by the formula:

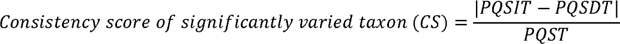

- PQSIT: project quantity with the significantly increased taxon
- PQSDT: project quantity with the significantly decreased taxon
- PQST: the total quantity of projects with the significantly varied taxon

The value of CS score (ranging from 0 to 1) indicates the consistency degree of the significant variation of the taxon across those comparisons with statistical differences. In addition, considering the consistency degree of the significant variation of the taxon among all comparisons, including those without significant difference, a total consistent variation score (TCS) is defined as follows:

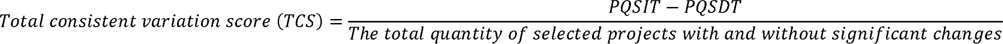

Thus, the TCS close to 1 or -1 indicates the higher degree of consistent increase or reduction across all the selected projects. The variation feature is visualized using consistency heatmap.

### Software development and implementation

OBMeta is implemented as a web server using JavaScript, HTML and cascading style sheets for fronted development. The core JavaScript library Vue.js (https://vuejs.org/) and Spring Boot (https://spring.io/projects/spring-boot) were used as the main frontend and backend frameworks respectively. The interactive communication was implemented using WebSocket. R in-house scripts were used for backend data processing, analyses, and visualizations. Open-source data management system MySQL (https://www.mysql.com/) is used for data persistent storage. OBMeta was deployed on Dell PowerEdge Server in the Children’s Hospital, Zhejiang University School of Medicine with 32 virtual CPUs (2.10Hz), 256 GB memory and 30T solid-state drives.

## Results

OBMeta is designed with three modules: (1) my project analysis, (2) cross-project validation, and (3) advanced validation. The overall workflow and analysis strategy are summarized in **Figure 1** and **Supplementary Figure 2**. The first module provides the comparative analysis including feature analysis, diversity analysis, functional analysis, and network analysis. The second module is to perform cross-project validation of gut microbial features across selected projects. The last module provides advanced validation for cross-phenotype or cross-intervention comparisons. All the results from main modules are available to download for further analysis and interpretation.

**Figure 1.**
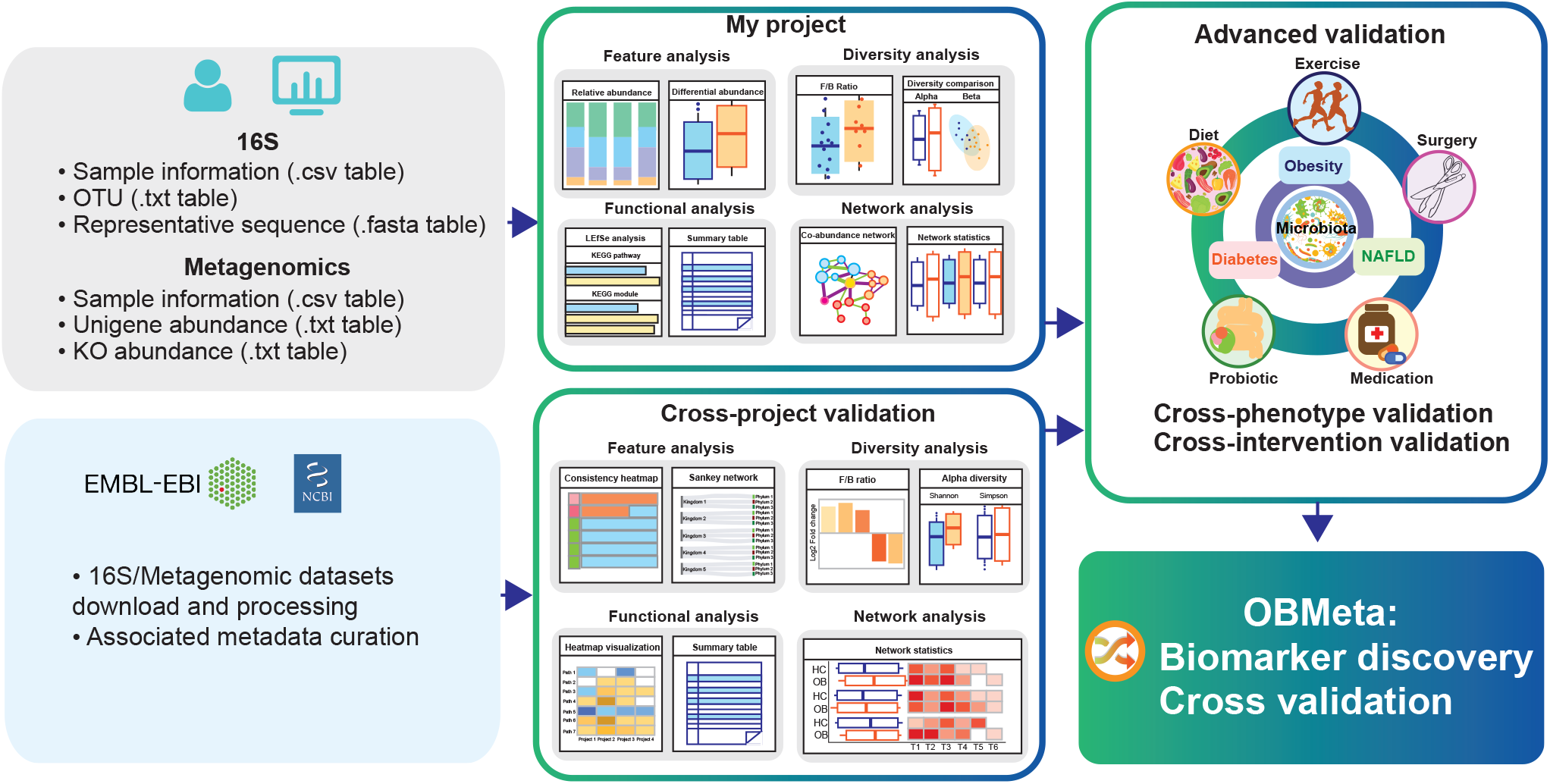
The workflow and functional features of OBMeta.

### My project analysis

#### Data uploading and filtering

16S rRNA sequencing and shotgun metagenomics are the most common sequencing strategies to characterize microbial compositions. OBMeta is designed to accept 16S rRNA marker gene data and metagenomics data. Two files are required for 16S data (a table with sample information and an OTU table) or metagenomics data (a table with sample information and a unigene table) to conduct comparative analysis. OBMeta also offers functional annotation and interpretation if representative sequence (16S rRNA data) or KO abundance (metagenomics data) datasets are available.

After uploading the data files, OBMeta will process the missing data automatically following the predefined steps: 1), to exclude features with a prevalence less than 20%; 2) to fill the zero with the 1/10 of minimum abundance; 3), to normalize data by DESeq2 using median of ratios method for 16S data. For metagenomics data input, OBMeta will produce gene abundance with TPM (transcripts per million) normalization using Salmon (version 1.2.1). This automatic filtering procedure can improve the statistical power and provide more robust results for downstream analysis.

#### Standardized analysis for local datasets

This module allows users to perform data analysis in four steps for their own datasets including feature analysis, diversity analysis, functional analysis and network analysis. In the feature analysis section, the OTU tables from marker gene or unigene can be assigned to different taxonomic level before conducting comparisons. The overview of microbial features is visualized as stacked bar plots (**Figure 2A**). OBMeta will adopt parametric or non-parametric univariate analysis based on data distribution to identify differential features displayed as box plots (**Supplementary Figure 3**) and summarized in a numerical table.

**Figure 2.**
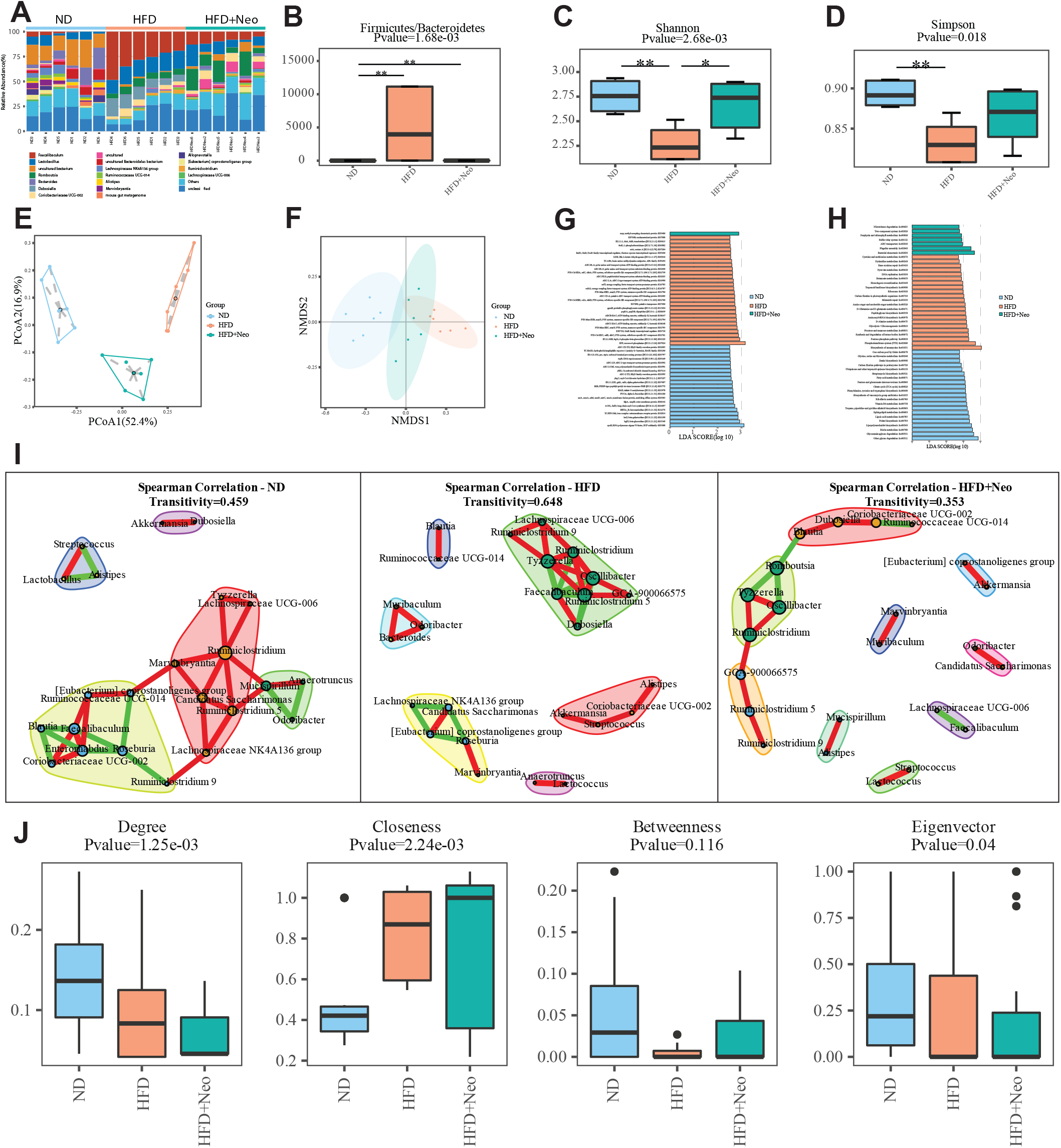
Outputs of the example datasets from ‘My project’ module of OBMeta. **(A)** A stacked bar plot of the relative abundance of gut microbiome at the genus level. **(B)** The Firmicutes/Bacteroidetes ratio of each group. **(C)** A bar plot of the Shannon index based on genus profile. **(D)** A bar plot of the Simpson index based on genus profile. **(E)** The ordination plot based on the Bray-Curtis distance at genus level using principal coordinate analysis (PCoA). **(F)** The ordination plot based on the Bray-Curtis distance at genus level using nonmetric multidimensional scaling (NMDS). **(G)** A bar plot of KO modules with significant LDA score (> 2) in LEfSe analysis (top 50 most significant modules). **(H)** A bar plot of functional pathways with significant LDA score (> 2) in LEfSe analysis (top 50 most significant pathways). **(I)** The co-abundance network from Spearman’s correlation analysis based on the top 30 most abundant genera for each group. **(J)** The box plots of the network properties including degree, closeness, betweenness, and eigenvector.

The Firmicutes/Bacteroidetes (F/B) ratio is widely considered as an important marker of intestinal homeostasis [29]. Increased F/B ratio is regarded as a potential signature in obesity [30]. So OBMeta offers the F/B ratio comparison among different groups (**Figure 2B**). In addition, the alpha-diversity is calculated at different taxonomic levels using Shannon index and Simpson index, which emphasizes the richness component and evenness component of diversity [31] (**Figure 2C and 2D**). The beta-diversity analysis provides two common distance measures (Bray-Curtis distance and Jaccard distance), which are represented by two-dimensional ordination plots based on principal coordinate analysis (PCoA) or non-metric multidimensional scaling (NMDS) (**Figure 2E and 2F**). The corresponding statistical significance is assessed using ANOSIM test (**Supplementary Figure 4**).

In functional analysis, OBMeta predicts functional profiles for 16S rRNA data or metagenomics data. Linear discriminant analysis (LDA) effect size (LEfSe) method has developed to identify robust features (KO modules and pathways) most likely to explain differences between groups by coupling non-parametric test for statistical significance with additional effect size estimation [32]. Log10 transformation of LDA cutoff > 2 is considered important as default. The top 50 differential modules or pathways are presented in bar plots (**Figure 2G and 2H**).

In the microbial community, ecological interactions between constituent taxa are vital in determining the overall structure and function of the community in host health and disease [33]. Also, the microbiome composition varies significantly between individuals raising the question whether there is a core microbiome that produces main effect with respect to the healthy and diseased states. To disclose the core composition in the associations of gut microbes, OBMeta is designed with network analysis module that helps users construct bacterial association network and calculate network topological properties for each group based on different taxonomic levels. The network nodes are colored according to the network modularity and the node sizes are proportionated with the degree (**Figure 2I**). The network properties (degree, closeness, betweenness, and eigenvectors) are compared using non-parametric test and presented in box plot (**Figure 2J**).

### Cross-project validation

#### Cross-project comparison and validation

The next main function of OBMeta is cross-project validation which allows users to validate the microbial features with multiple similar projects in the curated database. The curated database now includes 228 comparisons classified by phenotypes (case *vs.* control) and intervention strategies (case with intervention *vs.* case without intervention/ before intervention *vs.* after intervention) from a total of 88 projects in obesity, T2D, and NAFLD. Users can search similar projects according to the disease type, intervention strategy, and other filtering conditions including age, gender, species type, sample type and sequencing platform.

This module provides cross-project validation in four sections that are consistent with my project analysis. In feature analysis, OBMeta evaluates the consistent variation of taxa using CS and TCS, which are visualized in consistency heatmap (increase in orange and decrease in blue, **Figure 3A**), stacked bar plot (**Figure 3B**) and Sankey network (**Figure 3C**) at different taxonomic levels. The consistent features of all taxa are summarized in the table. In diversity analysis, OBMeta presents the cross-project comparison of the F/B ratio (**Figure 3D**) and alpha diversity (**Figure 3E**). In functional analysis, OBMeta provides a heatmap that summarize the pathway with significant LDA score in each project (**Figure 3F**). In network analysis, OBMeta calculates the network properties based on the top 30 most abundant taxa and presents the network property value (**Figure 3G**) and the mean correlation coefficient of each constituent member in heatmaps respectively. These multidimensional comparisons help efficiently identify the consistent microbial signatures with respect to comparative abundance, taxonomic diversity, functional pathway and bacterial interaction network for obesity-associated diseases.

**Figure 3.**
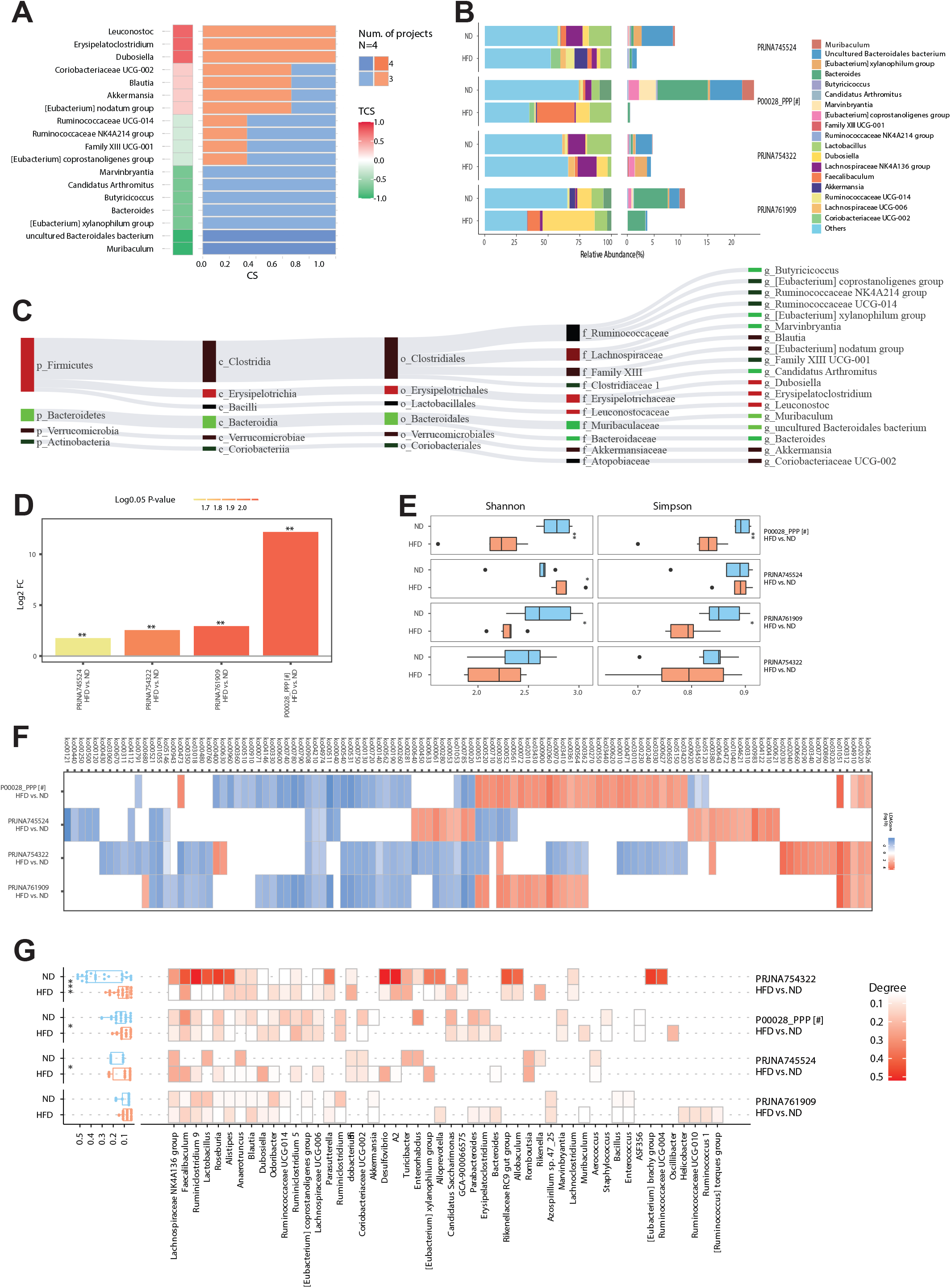
Outputs of the cross-project validation using the example comparison in ‘Phenotype’ classification. **(A)** Consistency heatmap of the significantly changed bacteria at the phylum level among the selected projects. The heatmap showed all phyla that significantly varied in at least 3 projects with the consistency score more than 0.6. **(B)** The stacked bar plot of the relative abundance of top 10 abundant genera and others (the left panel) and the consistently varied genera (the right panel). Local comparison was labeled with #. **(C)** Sankey network of the consistently varied genera that met the defaulted conditions. **(D)** The bar plot of the fold changes of the *Firmicutes*/*Bacteroidetes* ratio among different projects. It is coloured based on the log10 transformation of p-value in the comparison. **(E)** The box plots of alpha diversity (Shannon and Simpson indices based on the genus profile) in each group. **(F)** The heatmap of the significant pathways (LDA score > 2, LEfSe analysis) in any one of the selected projects. **(G)** The heatmap of the network property (degree) in each group based on the top 50 most abundant genera.

## Advanced validation

### Advanced validation via multi-disease and multi-intervention comparisons

Since obesity is closely associated with other metabolic diseases and various medical, nutritional, or physical interventions have been conducted targeting the gut microbiota, OBMeta offers multi-disease and multi-intervention evaluations of varied taxa in the module of advanced validation. The personal project and selected projects in cross-project validation are categorized as a whole by default to compare with other diseases (**Figure 4A**) or interventions (**Figure 4B**).

**Figure 4.**
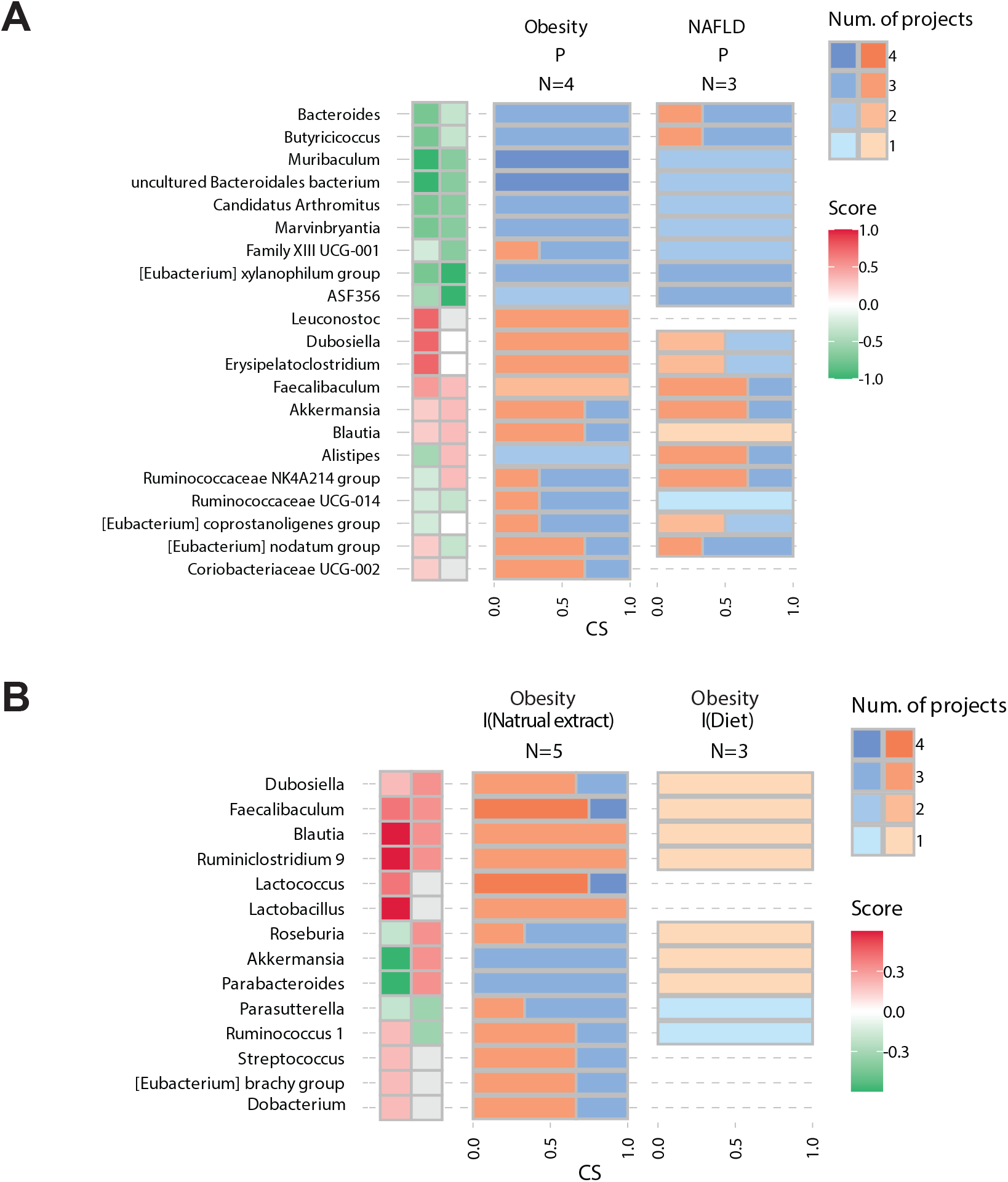
Example outputs of the advanced validation. **(A)** Consistency heatmap of the significant phyla between obesity and NAFLD. **(B)** Consistency heatmap of the significant phyla between the drug (natural extracts) and diet intervention with positive effects on obesity.

OBMeta is designed flexibly so that users can stop or restart their analysis in every module as they want. If users don’t have their own dataset for analysis and validation, they also can search on OBMeta based on the curated datasets for the microbial signature of interest.

## Case study

To illustrate the utility of OBMeta, we conducted a gut microbiome study on the beneficial effect of Neohesperidin (Neo, a natural polyphenol abundant in citrus fruits) in obesity using 8-week-old male wildtype (C57BL/6J) mice. Fecal samples were collected after 12 weeks on normal diet, high-fat diet and high-fat diet plus Neo (Project ID: PRJNA601832). This dataset includes three groups (label as ND, HFD and HFD + Neo). Two pairwise comparisons were conducted (HFD *vs.* ND and HFD + Neo *vs.* HFD) to validate the gut microbiome changes in obese mice with high-fat diet and effects of Neo intervention, respectively.

After file uploading, all the comparative analysis was conducted automatically in the ‘My project’ module. According to the results in the feature analysis section, we found that the genus *Faecalibaculum* significantly increased in the HFD group while Neo intervention can decrease its abundance to some extent (**Figure 2A**). The comparative analysis showed that *Leuconostoc*, *Faecalibaculum*, uncultured *bacterium*, *Dubosiella*, *Globicatella* and *Candidatus arthromitus* were the most significantly changed genera between groups (**Supplementary Figure 3**). In diversity analysis section, the *Firmicutes*/*Bacteroidetes* ratio was significantly increased in the HFD group regarding the increased *Firmicutes* and the almost depletion of *Bacteroidetes* (**Figure 2B**). In addition, the level of alpha diversity calculated by Shannon and Simpson indices obviously decreased in the HFD group compared to the ND and HFD + Neo groups (**Figure 2C and 2D**). The community structure also significantly varied according to the ordination plots (**Figure 2E and 2F**) and ANOSIM test (**Supplementary Figure 4**). In the functional analysis section, we found the significant KO modules (**Figure 2G**) and functional pathways (**Figure 2H**) in each group. In the network analysis section, the network plot indicates significant differences in the interactions between three groups (**Figure 2I**). For example, the beneficial genus *Akkermansia* positively correlated with *Dubosiella* in ND mice while it positively correlated with *Streptococcus* in HFD mice. The comparison of network properties also shows the degree and eigenvector was significantly decreased in HFD network which indicates less connections for the main genera, while the closeness was significantly increased in HFD network (**Figure 2J**).

Furthermore, to validate the significantly changes of this local comparison (HFD *vs.* ND), we selected similar projects (**Supplementary Table 2**) to identify the cross-project consistent variation. As **Figure 3A** shown, the genera *Muribaculum* and uncultured *Bacteroidales bacterium* were consistently decreased in high-fat-diet mice in totally 4 compared projects. Also, the genera *Leuconostoc*, *Eysipelatoclostridium* and *Dubosiella* were consistently increased, while *Eubacterium xylanophilum* group, *Bacteroides*, *Butyricicoccus*, *Candidatus arthromitus*, *Marvinbryantia*, and *Eubacterium coprostanoligenes* group consistently decreased in 3 of the selected projects. The stacked bar plot indicated the relative expressions of microbes specifically in each compared project (**Figure 3B**). We can observe that the relative abundance of the genera *Bacteroides* and uncultured *Bacteroides bacterium* with the most decrease in high-fat diet mice in local comparison (labeled with #) and PRJNA761909. The Sankey network demonstrated the consistently varied genera and their higher sources, which met the filtering conditions (project number = 3, CS = 0.6), indicating the increased abundance of phylum *Firmicutes* was accounted from the genera *Dubosiella*, *Erysipelatoclostridium* and *Leuconostoc* while the decrease of *Bacteroidetes* was mainly derived from the genera *Muribaculum*, uncultured *Bacteroidales bacterium* and *Bacteroides* in obesity (**Figure 3C**). In the diversity analysis, the *Firmicutes*/*Bacteroidetes* ratio showed consistently increased in selected projects (n = 4) (**Figure 3D**). Besides, local comparison shows significantly increased Shannon and Simpson indices, which was consistent with PRJNA761909 (**Figure 3E**). We also identified the pathways Ko00908 (zeatin biosynthesis), Ko04974 (protein digestion and absorption) and Ko04210 (apoptosis) are consistently decreased in 4 comparisons (**Figure 3F**). At last, the network analysis indicates that the degree of the genera was consistently increased in ND networks (**Figure 3G**). Collectively, *Muribaculum*, uncultured *Bacteroidales bacterium* and *Bacteroides* were potential biomarkers that decreased in high-fat-diet obesity. Besides the cross-project validation for HFD *vs.* ND comparison, we can also conduct validation for HFD + Neo *vs.* HFD comparison based on the similar projects with natural extracts intervention (**Supplementary Table 2**). The output format is same as above and users can explain based on their knowledge (**Supplementary Figure 5**). After cross-project validation, we can further put above findings to cross-phenotype comparison and cross-intervention comparison in ‘Advanced validation’ module. We selected projects of NAFLD using the same condition (**Supplementary Table 2**) to compare with HFD *vs.* ND comparison and found that the genera *Muribaculum* and uncultured *Bacteroidales bacterium* significantly decreased in obesity (n = 4) and NAFLD (n =1) (**Figure 4A**). Additionally, we can select projects of obesity with beneficial diet intervention to compare with the natural extracts intervention. The consistency heatmap (**Figure 4B**) indicates natural extracts and beneficial diets tend to consistently increase the abundance of *Blautia* and *Ruminiclostridium* 9.

## Discussion

OBMeta is developed with comprehensive capabilities including comparative metagenomics analysis, cross-project validation, cross-phenotype and cross-intervention comparisons. To the best of our knowledge, this is the first versatile tool comprising metagenomics data processing, comparative analysis, co-abundance network comparison, and multidimensional validation, which aims to identify microbial features and biomarkers of obesity-associated metabolic diseases.

With increasing high-quality public metagenomics datasets from studies of metabolic diseases, researchers have started to integrate independent public datasets/projects with the same phenotype or intervention to explore the consistent microbial signatures for validation. For example, several 16S rRNA sequencing studies were integrated to evaluate the microbial changes of beneficial diets (peach, wheat, quinoa, barley, cherry, raspberry, and apple) in healthy and obese animal models [34]. In addition, *Farnaz Fouladi* and his collogues have characterized robust and consistent microbial signatures across multiple projects with obese patients after Roux-en-Y Gastric bypass (RYGB) surgery using their own and existing sequencing data [35]. OBMeta can automatically conduct cross-project, cross-phenotype, and cross-intervention comparisons, making obesity-associated microbiome identification and validation more accurately and efficiently.

Over the past decade, several web-based metagenomics comparative tools have been developed to contribute the exploration of gut microbial features and biomarkers of different diseases. EBI-Metagenomics [36], VMPS [37] and MG-RAST [14] are developed primarily for raw sequence processing and annotation, they also provide limited statistical methods and visualizations. Busybee [17] is a new server for metagenomic data analysis in the form of assembled contigs or long reads. For single project exploration of comparative metagenomics, the METAGEN-assist [15] and Microbiome-Analyst [16] are representative tools comprising both univariate and multivariate statistical selections. Microbiome-Analyst also integrates with more newly developed statistical algorithms, functional annotation and visualization, and even allows users to upload multiple comparisons for potential biomarker identification in the version 2. Compared to Microbiome-Analyst, OBMeta evaluates the consistent or inconsistent alterations of gut microbial compositions and differential expressions in an efficient way by integrating public datasets of obesity-related metabolic diseases. Obviously, OBMeta is the first comparative metagenomics tool combining with comprehensive powerful function to compare, identify and validate microbial signature targeting on obesity-related diseases. More details of existing programs are summarized and compared in **Table 1**.

The limitations of the current version of OBMeta deserve to be mentioned. First, in addition to obesity, T2D and NAFLD, other obesity-associated diseases like hypertension and hyperlipidemia which could also be affected by gut microbiota were not included due to the limited number of public available studies so far. So far, we have included 4113 samples spanning 88 projects, however, the curated database may not cover all the associated datasets available in the public datasets. In the future, our team will continually update more well-designed projects and obesity-associated diseases to enlarge the curated database. It is also highly recommended that users may contribute their own projects or notify us any publicly available projects of interest through webserver link. Besides, more new statistical algorithms will be integrated into OBMeta for biological interpretation.

## Conclusion

OBMeta is the first comprehensive web-based tool to analyze and validate gut microbial signatures and biomarkers of obesity-associated metabolic diseases by streamlining the process of single-project analysis, cross-project validation, cross-phenotype and cross-intervention comparisons.

## Data availability

OBMeta is freely available at: http://obmeta.met-bioinformatics.cn/.

## Supplementary data

Supplementary data are available online at the time of publication.

## Authorship contribution

YN conceived and designed this project. CFX, YQG and JTH developed the web server. JTH, FZ and YN drafted and revised the manuscript. JTH and WXZ performed software testing. CFX, SYQ, JTH, and WXZ collected public datasets. FHL evaluated the quality of clinical and animal studies and public datasets included in OBMeta. GY supported the setup and maintenance of computational servers. CFX performed data processing and analysis.

## Acknowledgements

The authors acknowledged Ning Mei and Ye Wang from Zhejiang University School of Medicine on public datasets searching in this work.

## Funding

This work was supported by the National Key Research and Development Program of China [grant number 2021YFC2701904]; and the National Natural Science Foundation of China [grant number 82170583, 81900510].

## Key points

- User-friendly bioinformatics analysis for users’ local datasets, including feature analysis, diversity analysis, functional analysis and network analysis.
- Efficient cross-project identification and validation of obesity-associated microbial biomarkers between public datasets and the local dataset.
- Cross-phenotype and cross-intervention-based advanced validation to support preliminary findings.

## Notes

### Competing Interest Statement

The authors have declared no competing interest.

## References

1. Chooi YC, Ding C, Magkos F. The epidemiology of obesity, Metabolism 2019;92:6–10.

2. Franks PW, McCarthy MI. Exposing the exposures responsible for type 2 diabetes and obesity, Science 2016;354:69–73.

3. Riazi K, Azhari H, Charette JH et al. The prevalence and incidence of NAFLD worldwide: a systematic review and meta-analysis, Lancet Gastroenterol Hepatol 2022;7:851–861.

4. Khan SS, Ning H, Wilkins JT et al. Association of Body Mass Index With Lifetime Risk of Cardiovascular Disease and Compression of Morbidity, JAMA Cardiol 2018;3:280–287.

5. DeMarco VG, Aroor AR, Sowers JR. The pathophysiology of hypertension in patients with obesity, Nat Rev Endocrinol 2014;10:364–376.

6. Jandhyala SM, Talukdar R, Subramanyam C et al. Role of the normal gut microbiota, World J Gastroenterol 2015;21:8787–8803.

7. Qin J, Li Y, Cai Z et al. A metagenome-wide association study of gut microbiota in type 2 diabetes, Nature 2012;490:55–60.

8. Boulange CL, Neves AL, Chilloux J et al. Impact of the gut microbiota on inflammation, obesity, and metabolic disease, Genome Med 2016;8:42.

9. Tolhurst G, Heffron H, Lam YS et al. Short-chain fatty acids stimulate glucagon-like peptide-1 secretion via the G-protein-coupled receptor FFAR2, Diabetes 2012;61:364–371.

10. Marques FZ, Mackay CR, Kaye DM. Beyond gut feelings: how the gut microbiota regulates blood pressure, Nat Rev Cardiol 2018;15:20–32.

11. Wang J, Tang H, Zhang C et al. Modulation of gut microbiota during probiotic-mediated attenuation of metabolic syndrome in high fat diet-fed mice, ISME J 2015;9:1–15.

12. Tomova A, Bukovsky I, Rembert E et al. The Effects of Vegetarian and Vegan Diets on Gut Microbiota, Front Nutr 2019;6:47.

13. Ulker I, Yildiran H. The effects of bariatric surgery on gut microbiota in patients with obesity: a review of the literature, Biosci Microbiota Food Health 2019;38:3–9.

14. Wilke A, Bischof J, Gerlach W et al. The MG-RAST metagenomics database and portal in 2015, Nucleic Acids Res 2016;44:D590–594.

15. Arndt D, Xia J, Liu Y et al. METAGENassist: a comprehensive web server for comparative metagenomics, Nucleic Acids Res 2012;40:W88–95.

16. Dhariwal A, Chong J, Habib S et al. MicrobiomeAnalyst: a web-based tool for comprehensive statistical, visual and meta-analysis of microbiome data, Nucleic Acids Res 2017;45:W180–W188.

17. Schmartz GP, Hirsch P, Amand J et al. BusyBee Web: towards comprehensive and differential composition-based metagenomic binning, Nucleic Acids Res 2022.

18. Kodama Y, Shumway M, Leinonen R et al. The Sequence Read Archive: explosive growth of sequencing data, Nucleic Acids Res 2012;40:D54–56.

19. Harrison PW, Alako B, Amid C et al. The European Nucleotide Archive in 2018, Nucleic Acids Res 2019;47:D84–D88.

20. Hall M, Beiko RG. 16S rRNA Gene Analysis with QIIME2, Methods Mol Biol 2018;1849:113–129.

21. Prodan A, Tremaroli V, Brolin H et al. Comparing bioinformatic pipelines for microbial 16S rRNA amplicon sequencing, PLoS One 2020;15:e0227434.

22. Douglas GM, Maffei VJ, Zaneveld JR et al. PICRUSt2 for prediction of metagenome functions, Nat Biotechnol 2020;38:685–688.

23. Bolger AM, Lohse M, Usadel B. Trimmomatic: a flexible trimmer for Illumina sequence data, Bioinformatics 2014;30:2114–2120.

24. Langmead B, Salzberg SL. Fast gapped-read alignment with Bowtie 2, Nat Methods 2012;9:357–359.

25. Li D, Luo R, Liu CM et al. MEGAHIT v1.0: A fast and scalable metagenome assembler driven by advanced methodologies and community practices, Methods 2016;102:3–11.

26. Gurevich A, Saveliev V, Vyahhi N et al. QUAST: quality assessment tool for genome assemblies, Bioinformatics 2013;29:1072–1075.

27. Hyatt D, LoCascio PF, Hauser LJ et al. Gene and translation initiation site prediction in metagenomic sequences, Bioinformatics 2012;28:2223–2230.

28. Patro R, Duggal G, Love MI et al. Salmon provides fast and bias-aware quantification of transcript expression, Nat Methods 2017;14:417–419.

29. Stojanov S, Berlec A, Strukelj B. The Influence of Probiotics on the Firmicutes/Bacteroidetes Ratio in the Treatment of Obesity and Inflammatory Bowel disease, Microorganisms 2020;8.

30. Magne F, Gotteland M, Gauthier L et al. The Firmicutes/Bacteroidetes Ratio: A Relevant Marker of Gut Dysbiosis in Obese Patients?, Nutrients 2020;12.

31. DeJong TM. A Comparison of Three Diversity Indices Based on Their Components of Richness and Evenness, Oikos 1975;26:222–227.

32. Segata N, Izard J, Waldron L et al. Metagenomic biomarker discovery and explanation, Genome Biol 2011;12:R60.

33. Loftus M, Hassouneh SA, Yooseph S. Bacterial associations in the healthy human gut microbiome across populations, Sci Rep 2021;11:2828.

34. Garcia-Mazcorro JF, Kawas JR, Licona Cassani C et al. Different analysis strategies of 16S rRNA gene data from rodent studies generate contrasting views of gut bacterial communities associated with diet, health and obesity, PeerJ 2020;8:e10372.

35. Fouladi F, Carroll IM, Sharpton TJ et al. A microbial signature following bariatric surgery is robustly consistent across multiple cohorts, Gut Microbes 2021;13:1930872.

36. Hunter S, Corbett M, Denise H et al. EBI metagenomics--a new resource for the analysis and archiving of metagenomic data, Nucleic Acids Res 2014;42:D600–606.

37. Huse SM, Mark Welch DB, Voorhis A et al. VAMPS: a website for visualization and analysis of microbial population structures, BMC Bioinformatics 2014;15:41.

