## Supplementary Data for "OBMeta: a comprehensive web server to analyze and validate gut microbial features and biomarkers for obesity-associated metabolic diseases"

Additional material

**1. Supplementary table**

**1.1 Supplementary Table 1** **The software summary used in data processing**

| **Name** | **Version** | **Parameter** |
| --- | --- | --- |
| FastQC | 0.11.8 | Default |
| QIIME 2 | 2020.2 | Default |
| DADA 2 | 1.16.0 | Default |
| PICRUSt2 | 2.5.0 | Default |
| Trimmomatic | 0.39 | ILLUMINACLIP:TruSeq2-PE.fa:2:40:15  SLIDINGWINDOW:6:20 LEADING:3  TRAILING:3 MINLEN:50 |
| Bowtie 2 | 2.4.1 | Default |
| MEGAHIT | 1.2.9 | --min-contig-len 500 --min-count 2 |
| MetaGeneMark | 3.38 | -a -d -f G -m |
| CD-HIT | 4.8.1 | -c 0.95 -aS 0.9 -g 1 -d 0 -M 0 |
| Diamond | 0.9.34 | -e 1e-5 |
| Salmon | 1.2.1 | Default |

**1.2 Supplementary table 2 The projects used for cross-project validation and advanced validation**

| **Filtering condition** | **Cross-project validation** | | **Advanced validation** | |
| --- | --- | --- | --- | --- |
| **Disease type** | Obesity | Obesity | NAFLD | Obesity |
| **Classification** | Phenotype | Intervention | Phenotype | Intervention |
| **Age stage** | Adult | Adult | Adult | Adult |
| **Gender** | Male | Male | Male | Male |
| **Species type** | Mus musculus | Mus musculus | Mus musculus | Mus musculus |
| **Sample type** | Feces | Feces | Feces | Feces |
| **Sequencing platform** | Illumina Miseq | Illumina Miseq | Illumina Miseq | Illumina Miseq |
| **Intervention type** | - | Drug | - | Diet |
| **Intervention effect** | - | Positive | - | Positive |
| **Project** | PRJNA745524, PRJNA761909, PRJNA761909 | PRJNA760837, PRJNA761909, PRJNA811511, PRJDB9484, | PRJEB36797, PRJEB48939, PRJDB7523 | PRJNA745524,  PRJNA281761,  PRJEB36801 |

**2. Supplementary Figure**

**2.1 Supplementary Figure 1**

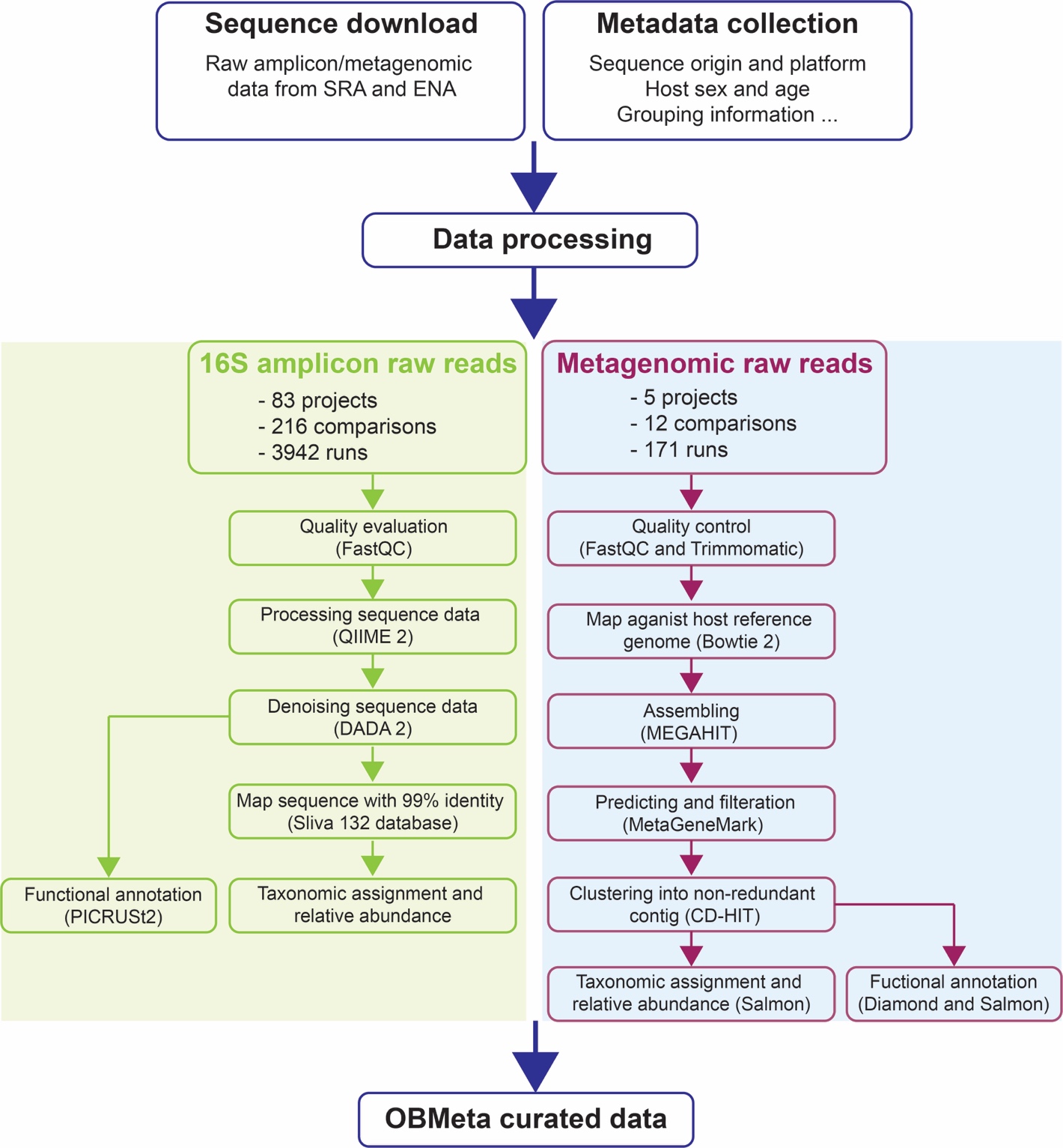

**Supplementary Figure 1.** Overall workflow of data processing for the curated data of OBMeta.

**2.2 Supplementary Figure 2**

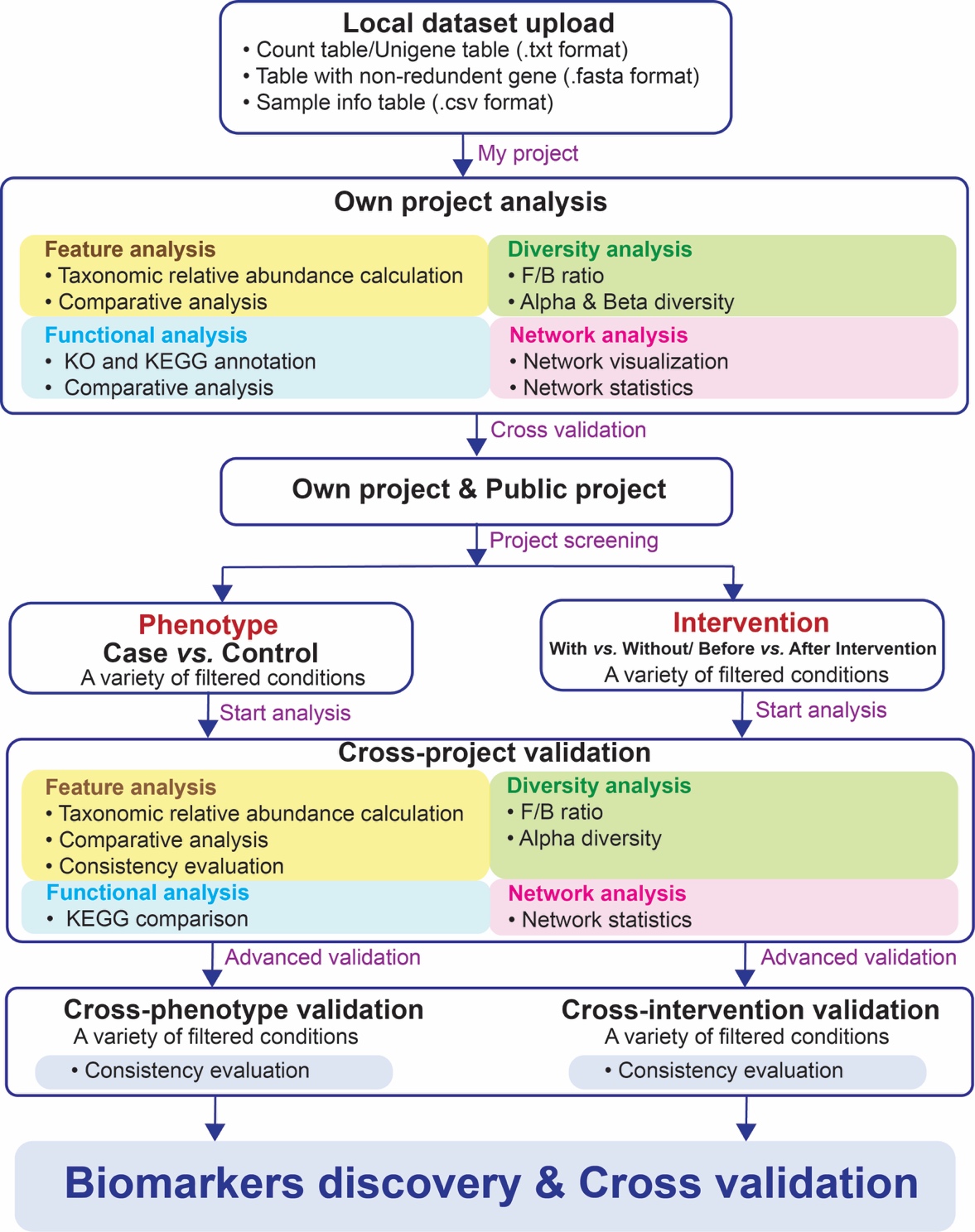

**Supplementary Figure 2.** The recommended analysis strategy using OBMeta.

**2.3 Supplementary Figure 3**

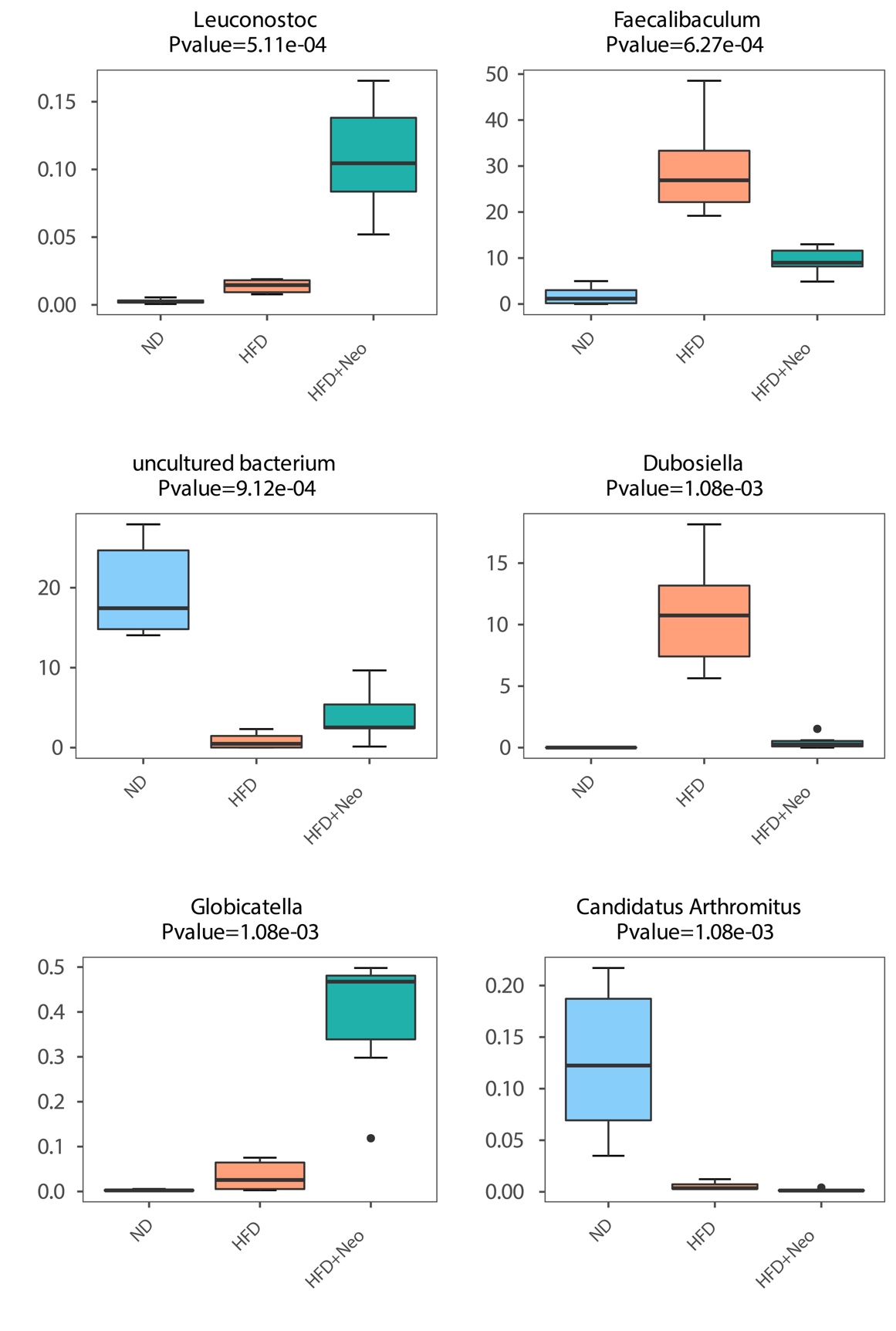

**Supplementary Figure 3.** One example output of the local dataset in feature analysis (at genus level). All phyla will be compared in OBMeta and visualized in box plots in download file. In the real-time interface, OBMeta only presents top 6 most differentially abundant phyla.

**2.4 Supplementary Figure 4**

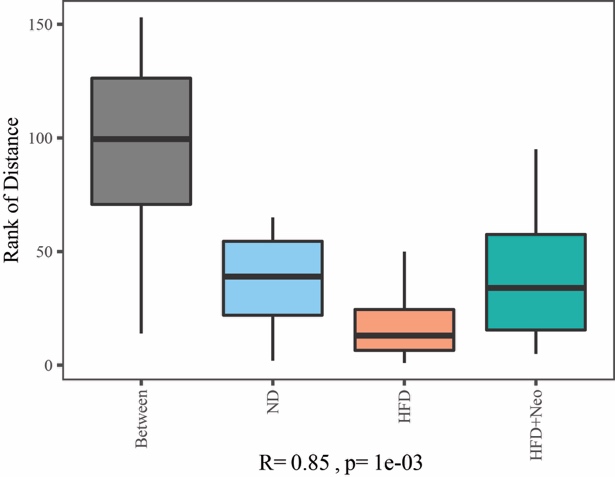

**Supplementary Figure 4.** Analysis of similarities (ANOSIM) test used in beta diversity comparison. The box plot shows the ranked dissimilarities between groups and the ranked dissimilarities within groups.

**2.5 Supplementary Figure 5**

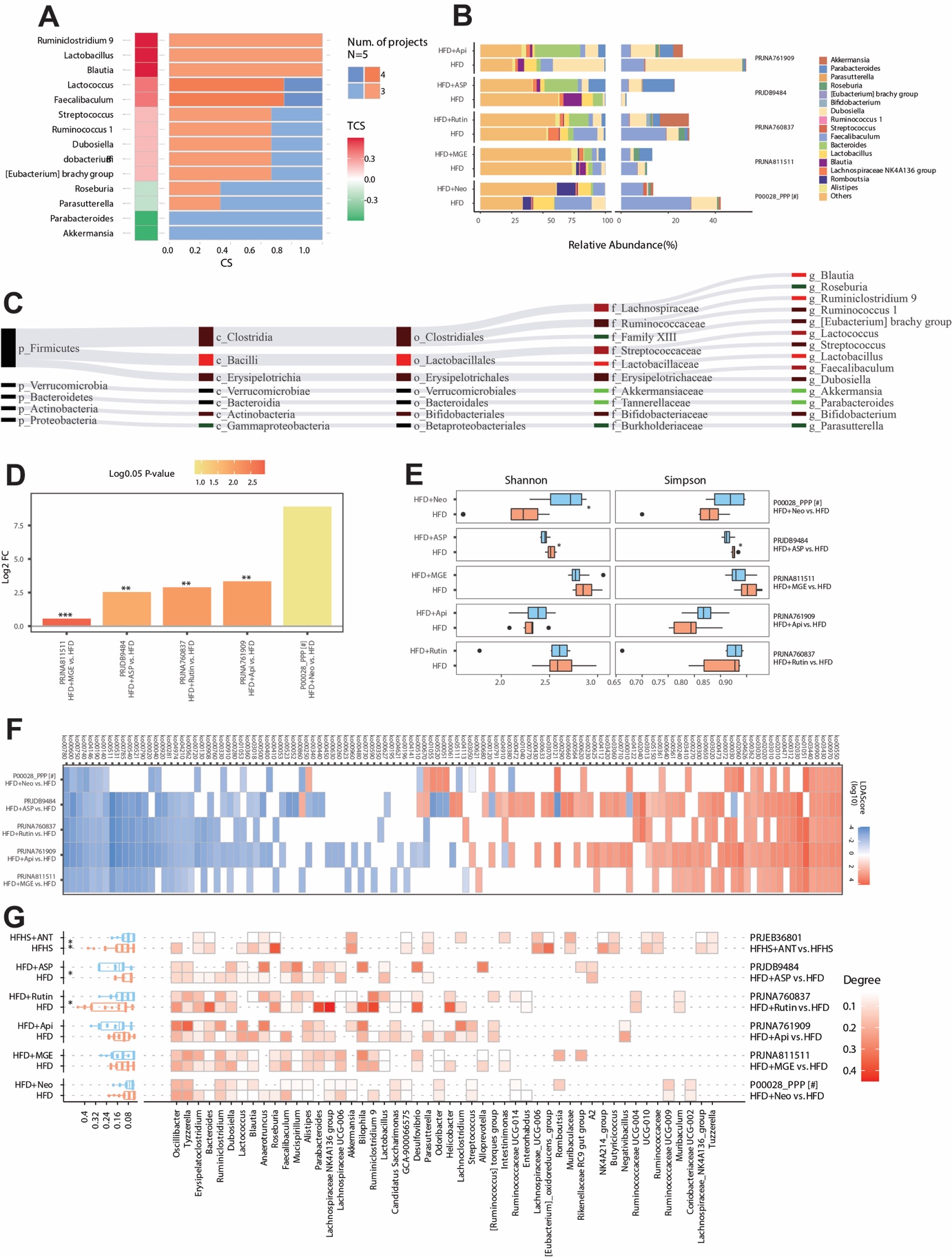

**Supplementary Figure 5.** Example outputs of the cross-project validation in ‘Intervention’ classification. **(A)** The consistency heatmap evaluated the intervention effect on gut microbiome at the genus level through multiple projects using default settings (project number = 3, consistency score = 0.6). **(B)** Sankey network of the consistently varied genus in selected projects. **(C)** The stacked bar plot of the relative abundance of all genera (the left panel) and the consistently varied genera (the right panel). **(D)** The log2 transformation of the fold change of the *Firmicutes*/*Bacteroidetes* ratio. **(E)** Alpha diversity. It is measured by Shannon and Simpson indices using the genus profile. **(F)** The functional pathway with significant LDA score based on LEfSe analysis. **(G)** The comparison of the network property (degree) in each group. The network is constructed based on the top 30 most abundant genera. The heatmap shows the degree value of each genus in the network.
